# Statistics in service of metascience: Measuring replication distance with reproducibility rate

**DOI:** 10.1101/2024.08.05.606644

**Authors:** Erkan Buzbas, Berna Devezer

## Abstract

Motivated by the recent putative reproducibility crisis, we discuss the relationship between replicability of scientific studies, reproducibility of results obtained in these replications, and the philosophy of statistics. Our approach focuses on challenges in specifying scientific studies for scientific inference via statistical inference, and is complementary to classical discussions in philosophy of statistics. We particularly consider the challenges in replicating studies exactly, using the notion of the idealized experiment. We argue against treating reproducibility as an inherently desirable property of scientific results, and in favor of viewing it as a tool to measure distance between an original study and its replications. To sensibly study the implications of replicability and results reproducibility on inference, such a measure of replication distance is needed. We present an effort to delineate such a framework here, addressing some challenges in capturing the components of scientific studies while identifying others as ongoing issues. We illustrate our measure of replication distance by simulations using a toy example. Rather than replications, we present purposefully planned modifications as an appropriate tool to inform scientific inquiry. Our ability to measure replication distance serves scientists in their search for replication-ready studies. We believe that likelihood-based and evidential approaches may play a critical role towards building a statistics that effectively serves the practical needs of science.

## 1 Introduction

Discussions on the future of scientific inference and its relation to statistical inference are rare and challenging. As the sciences progress, the need for a broader philosophy that forges deeper connections between science and statistics becomes evident, heightening the demand for a unified statistical philosophy. We are excited to contribute to this intriguing special issue and offer our metascientific perspective on some timely questions.

Our tone is philosophical, stemming from our theoretical research program on statistical theory in and of metascience, and we hope it complements Taper et al. (2022). We raise a number of questions about improving scientific inference via statistical inference. We do not (yet) have all the answers, nonetheless we make some theoretical progress in this paper advancing a measure of distance between an original experiment and its replication that uses reproducibility rate as a metric. We present this measure to illustrate our perspective and chart one path toward closing the gap between statistical theory and scientific inference.

Our central argument is that a robust philosophy of statistics cannot thrive in isolation, detached from the fundamental challenges involving some meta-notions that have traditionally been viewed as beyond its scope^1^. By *meta-notions*, we mean concepts that are invoked by scientific inference and may have a bearing on statistical inference and beyond. Here we specifically focus on the relationship between replicability of scientific studies, reproducibility of results obtained in these replications, and the philosophy of statistics. We use replicability in the sense of repeatability of a scientific study and reproducibility in the sense of obtaining the same statistical result (conditional on a result type) in a sequence of replicated studies^2^. By philosophy of statistics we mean the approaches of statistical schools to inference and the theoretical frameworks advanced by these schools.

If a mature and coherent philosophy of statistics is to formally incorporate important meta-notions, an expansion of theoretical frameworks in statistics seems needed. For example, in the context of replicability and results reproducibility, we believe that a necessary first step is to develop a well-defined distance between scientific studies. The reason is straightforward. Statistical theory already allows us to appraise and study the outcomes of exact replication studies. However, that hardly helps us interpret the results of replications in practice since exact replication is an ideal that only exists in theory. Real-life replications are subject to many kinds of deviations from an original study. Statistics has been largely silent on how we should adjust our appraisals of results reproducibility accordingly. An attempt to delineate replication distance is provided in this paper with some progress, albeit not a fully satisfactory one, due to challenges associated in formalizing some components of scientific studies.

Contrary to a popular stance in the putative reproducibility crisis, we do not view results reproducibility as an inherently desirable property of studies. Instead, we consider it a valuable tool in assessing the quantitative impact of variation between studies. Hence, we focus on a distance between studies based on reproducibility rates of results. We advance the idea of purposeful modifications of studies and describe the notion of Minimum Viable Experiment to replicate (MVE) (Devezer and Buzbas, 2022) as a reference point to assess replication-readiness of studies. Since our approach is complementary to classical discussions on philosophy of statistics, we assume some background knowledge on the part of the reader. Succinct reviews can be found in Durbin (1987); Lindley (2000); Taper and Ponciano (2016) and would be helpful to make sense of the perspective taken in this paper.

Quantitative sciences have profoundly shifted their approach to scientific inference over recent decades, particularly with the help of substantial advancements in computing power. It is remarkable that the gain is computational but the resulting shifts are also conceptual. Computation has made previously not pursued ideas practically possible. Examples include a growing emphasis on: 1) predictive inference using learning-based approaches that have been fundamental in the refinement of machine learning and artificial intelligence applications, 2) causal inference incorporating logical pathways as a fundamental aspect of statistical models to establish dependencies and infer causality in regression models, 3) simulation-based computational methods to perform inference under large models often characterized by stochastic processes, 4) metascientific methods to improve the replicability of and results reproducibility in scientific studies. This short list of examples points to a shift toward addressing a broad range of problems with novel approaches to inference. Examples (1) and (2) are about inferential frameworks: Predictive inference judges the adequacy of a model by its predictive success and is willing to accept a model as a black-box as long as it is successful in predicting future observable outcomes. Causal inference highly values a detailed understanding of the mechanism that generates the data over predictive accuracy. Example (3) involves tools to approach inference. Simulation-based methods investigate generic algorithmic approaches to computer intensive statistics, irrespective of the modeling context. Example (4) concerns how data are generated in scientific studies. Replicability of scientific studies and results reproducibility testify to the degree of control exercised in gaining the information targeted and the standardization of its inferential implications.

Differences in goals aside, a common driver behind all of these perspective shifts is the necessity for statistics to serve the practical needs of science rather than being driven solely by philosophically motivated inquiries. Frequentists, pure likelihoodists, and Bayesians alike have contributed solutions toward advancing statistical inference in various domains. This *statistics in service of science* position has consequences for developing a sound philosophy of statistics. For one, statistics cannot be easily separated from practical considerations even in its philosophy. In Lindley’s words: “Only as theoreticians can we –statisticians– exist alone. Even there we suffer if we remain too divorced from the science”(Lindley, 2000, p.294). Further, statistics needs to stay relevant as the machinery of scientific inference under uncertainty while *simultaneously* growing in a principled way. This requires building a philosophy of statistics that is compatible with the practice of rapid perspective changes in science. As Box (1976) notes, rapid changes require flexibility to profit from the confrontations of practice and theory. To achieve such flexibility, it seems that statistics needs to expand its limited operating mode of “assume a model, develop a method, and perform inference for unknowns” by incorporating fundamental *meta-notions* formally into its theory. These meta-notions include: interpretations of probability (i.e., symmetry-based, propensity, frequency, belief-based), nature of evidence, practicability of replicability of studies and results reproducibility, scale of measurements, decision, causality, open-world/closed-world assumptions, and simulation. Some of this is already done. For example, we have a good understanding of frequentist, pure-likelihood, and a variety of Bayesian approaches based on different interpretations of probability. Causality, decision, and simulation are also well-studied. However, other notions such as the nature of evidence (where evidential approach to inference can help), practicability of replicability of studies and results reproducibility, scales of measurement of variables, and open-world/closed-world assumptions have not been prioritized in statistical theory. Incorporating all these meta-notions in a coherent inferential framework explicitly seems to be a daunting task. Indeed, discussions on meta-notions regularly rekindle old philosophical debates in statistics (e.g., Birnbaum’s likelihood principle, (Birnbaum, 1962) and discussions in and on Mayo (2014)). These debates jump out like monsters hiding under the bed from time to time in statistics literature with seemingly little progress. The task of incorporating them formally into a widely acceptable philosophy of statistics then seems doubly challenging.

Per our expertise and to keep the ideas focused, here we look at aspects of scientific inference via statistical inference that are only relevant to replicability of scientific studies and results reproducibility. Our thoughts are organized as follows. In the next section, we give some background on the problem known as reproducibility crisis. We emphasize the missing relationship between theoretical statistics and reality of many scientific studies, and highlight some misinterpretations of the notion of replicability and results reproducibility. We then review some previous theoretical results on results reproducibility, based on the operational notion of *idealized experiment*. We emphasize the need for a distance between studies to study results reproducibility properly and introduce one such distance based on reproducibility rates as a fundamental currency. We provide an illustration of our measure of replication distance, presenting an analysis of simulated data under a toy example. We also introduce the idea of purposefully modified studies and discuss the notion of the Minimum Viable Experiment to replicate (MVE) as an alternative to replications. We conclude by reflecting on how our ideas connect to evidential theory.

## 2 Background on replications and results reproducibility

The problem of nonreproducibility of major scientific claims in some fields has come to the forefront in the last decade, followed by what is dubbed as a reproducibility or replication crisis, and the rise of metascience as a field. This putative crisis typically refers to instances where published research results have failed to be corroborated by a sequence of scientific studies. While this state of affairs has largely been considered a crisis of practice, we believe it also points to a crisis of conceptual understanding (Buzbas et al., 2023). Replication and reproducibility are methodological concepts, and therefore they can be formally studied to build a strong, internally consistent, verifiable theoretical foundation of metascience. Such a theoretical foundation^3^ aids our understanding of the drivers of nonreproducibility and help inform future scientific practice and inference.

A small but highly successful literature has begun to emerge using models to study results reproducibility (e.g., Bak-Coleman et al., 2022; Bonifay et al., 2024; Davis-Stober et al., 2024; Devezer et al., 2019; Fanelli, 2020; Fanelli et al., 2022; McElreath and Smaldino, 2015; Nissen et al., 2016; Smaldino and McElreath, 2016). This literature demonstrates the potential for formal tools to illuminate a more general theory of connections between scientific inference and statistical inference. Formalizing these connections requires formalization of scientific studies, their replications, and reproducibility of results from these replications. The topic has attracted attention from philosophers and scientists, but perhaps unsurprisingly not as much from theoretical statisticians. Reconciling scientific inference with statistical inference theoretically requires ideas from statistics, philosophy, and social sciences alike. Some fundamental problems on the connections between scientific inference and statistical inference include: (i) how to incorporate background information and methods preceding data collection (pre-data methods, hereafter) such as choice of topic, sampling procedures, measures, and treatments into the assumptions of a statistical model, (ii) how to formally translate a scientific hypothesis into a statistical hypothesis, and (iii) how to compare results from various post-data methods. Formal solutions to these problems will necessarily have an impact on a satisfactory philosophical approach to statistics that we are willing to adopt.

### 2.1 Replication as an obscure object

Statisticians and scientists have not communicated well on what replication studies mean. Statistics has been a methodology-centric field, and statisticians have paid considerable attention to the performance of statistical methods with only a cursory nod to meta-notions regarding replications and reproducibility. Once two studies assume the same model, to a statistician the difference between these studies technically disappear, yielding random samples drawn from the same population. Scientists who have conducted real-life experiments would (hopefully) acknowledge that this is an overly ambitious assumption. Nevertheless, statisticians have been relatively conservative in incorporating components of real-life scientific studies when developing inferential methods. A common approach in statistical method development is to assume a model for the data generating mechanism and move to investigating the properties of newly devised methods under this model (presumably to establish superiority to an existing method). Statistics rarely asks how to satisfy the model assumptions in practice. This problem is believed to lie within the domain of the subject matter from which the data are derived. We believe this approach is an impediment for scientific and statistical progress alike. It is one thing to refer to a hypothetical sequence of independent tosses of a perfectly symmetric coin given the parameters and deduce that the number of heads is Binomially distributed, and another thing to refer to a sequence of real-life studies performed at different labs, times, designs, populations. Assuming that these two sequences are qualitatively equivalent to facilitate real-life scientific inference is wishful thinking at best. Statistical methods often reference idealized sequences, such as coin tosses, and tend to avoid the complexities of real-life sequences. Similar problems arise when statistical analyses are performed under modeling assumptions chosen for their mathematical convenience because methods to perform inference under these assumptions are readily available. A classical example is conjugate models in Bayesian statistics.

One of the early approaches taken to reconcile results from non-exact replication studies was meta-analytic efforts to reconcile effect sizes from multiple studies (see the recent review by Bogomolov and Heller (2023)). Recent applications include systematic replication studies in social sciences where concerns of irreproducibility came to the forefront in the last decade (Klein et al., 2022, 2018) and their comparisons to meta-analytic results (Kvarven et al., 2020). Further, random effects model formulations for non-exact replications McShane et al. (2022) or even metastudies themselves have been suggested (Baribault et al., 2018). These and other approaches contribute to our understanding of results from non-exact replications. However, these efforts are based on the traditional statistical approach aforementioned in the preceding paragraph, which we think has limitations in understanding non-exact replications and results obtained from them. Continuing with these two examples, meta-analytic methods are not well-equipped to address differences between an original study and its replications arising from pre-data methods such as differences in data types, measurement scales, or data entry errors reflected in data structure. They do well what they aim at–combining results from different studies–but even this comes with additional assumptions about the methods they apply. They are simply not designed (and they do not claim) to accommodate all aspects of studies that might affect reproducibility rates. Random effects models accommodate individual or subpopulation level variability in the data generating mechanism, again under additional assumptions (e.g., expectation zero, normal distribution of random effects), but not methods (pre- or post-data). We believe that a more fundamental framework for formulating meta-notions involved in designing and performing studies from first principles, and incorporating these into study of replications holistically is needed to make substantial progress.

On the upside, mathematical issues arising from sample properties have been studied extensively within applied contexts. Examples include dependent observations, missing data, and hierarchical models. Unfortunately, the drivers of these properties have not received as much attention. Factors such as the background information of scientists in designing and performing a study and pre-data methods chosen in the implementation of a study have not been core considerations in statistical inferential approaches. Systematic investigations into which assumptions of an original study have been met or need to be met in a replication study are virtually non-existent. Typically, when designing a replication study, researchers make (at most) subjective judgments regarding the similarity or closeness to the original experiment. Sometimes differences are introduced not due to changing conditions or resource constraints, but as proposed improvements (e.g., “larger sample size is better”). Sometimes designs and methodologies diverge arbitrarily due to sampling different populations, using different measures, and employing different modes of experimental administration (see Buzbas and Devezer, 2023, for a case example). These differences affect the inferences drawn from the replication study and obscure the evidence supporting or refuting the original result. If a result fails to reproduce, is it because an original positive association was spurious or because of the variations in the study design and implementation? Methods to quantify the distance between studies and to connect that distance to the evidential value of data for a given scientific question would improve learning from replications. We believe it should primarily be the statisticians’–not the scientists’–responsibility to formalize (i.e., mathematically) the assumptions of scientific studies and properly incorporate them into statistical models and methods.

### 2.2 Reproducibility as a misunderstood concept

There are some misconceptions about what the reproducibility rate of a given result means with respect to the truth value of that result. For example, various versions of the following two statements are often reiterated in the context of reproducibility crisis: (i) A result that has high reproducibility rate implies that it is (likely to be) true (i.e., a true scientific discovery) or equivalently, a result that has low reproducibility rate implies that it is (likely to be) false (i.e., not a scientific discovery). (ii) A true result implies high reproducibility rate (or equivalently, a false result implies a low reproducibility rate). Both statements are false (and trivially so) (see Devezer et al., 2019, for the complete formal argument regarding the validity of these statements). The key to correct interpretation of results reproducibility is that a true reproducibility rate of a result is a parameter of a study but the truth value of the result is not the only determining factor of this parameter in a sequence of exact or non-exact replication studies.

For example, to see that (i) is false, assume that the goal is to estimate a continuous parameter *µ* ∈ ℝ of a model. For a sequence of studies, choose the method: return 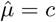 always, where *c* is a constant. The reproducibility rate of 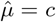 is 1 irrespective of data but the result is false with probability 1 for *µ≠ c*. Similarly, to see that (ii) is false, assume the true model generating the data has high noise to signal ratio with insufficient information to detect the signal. In the context of hypothesis testing of a null hypothesis with frequentist approach, consider a low powered statistical test. Assume that the null hypothesis is false. The reproducibility rate of rejecting the null hypothesis will be low but nonetheless the alternative hypothesis is true. These examples simply illustrate how easy it is to shift the reproducibility rate by changing the parameters of a study. We invite unconvinced readers to read our prior work cited here to find many mundane examples of the myriad ways this is achieved in practice, not limited to the examples above.

These misconceptions stem from the conflation of the *truth of a result* with the *reproducibility of a result*. The truth of a result depends on the statistical properties of the probabilistic mechanism generating the data. The reproducibility rate, on the other hand, also depends on the properties of the assumed models, methods, and the data structure of a study as well as the differences of the replication from the original study. In practice, even well-designed studies using statistical methods with well-established properties might yield different reproducibility rates for a result. This becomes clear, for example, if we consider different results obtained by different legitimate approaches to statistical inference given the same data. Akaike’s Information Criterion, Schwarz Information Criterion, posterior probability of a model, and adjusted *R*^(2)^ have all reasonable heuristic justifications (as well as some disadvantages) for the model selection problem. However, they will not always select the *same model* in unison –let alone the *true model*– in a given set of models, leading to highly variable reproducibility rates (see Devezer et al., 2019, for a detailed simulation example). Even before applying statistics on the data, differences in pre-data methods are also sufficient to show that there is a tenuous relationship between the truth value of a result and its reproducibility rate considering how many different studies can be conceived to answer the same scientific question (see Gelman and Loken, 2013). In practice, reproducibility rate speaks more closely to the design and components of original and replication studies than to the true data generating mechanism.

### 2.3 Reproducibility rate as a distance between a study and its replication

Based on the picture presented thus far, we conclude that the value of the notion of results reproducibility from replications in improving scientific inference via statistical inference does not lie in its mapping on the truth value of a result. Replications are conditional on an original study, which aims to sample a distribution of interest. Results reproducibility is with respect to a particular result type that is evaluated using a sample conditional on this distribution of interest. Replications and results reproducibility, therefore, are subject to epistemic inquiry in isolation, without reference to truth. Isolation from truth of a result to study replications and results reproducibility is a substantial technical gain. The truth is unknown and developing methods with reference to it is fraught. This idea of *isolation from truth* is also well-aligned with dropping the reference to true model generating the data in evidential theory of statistics (Lele, 2014).

We believe that a meaningful use of the notion of reproducibility–specifically the reproducibility rate of a result–is as a *currency* in quantifying statistical differences between scientific studies. As a first step in this quantification, a scientific study needs to be formalized with clearly defined components. We refer to this notion as an *idealized experiment*, which was introduced in Devezer et al. (2019) and formalized in Buzbas et al. (2023). An idealized experiment is a mathematical object that defines a scientific study in terms of background knowledge, pre-data methods, statistical model, post-data methods (i.e., statistical analysis), and data (structure and values). Conditional on a well-defined result type that is the goal of a study, exact replications of an idealized experiment yield a true reproducibility rate for that experiment. *Experiment* here is used as a convenience to represent any scientific study whose results are uncertain and the population is conducive to probability sampling, including but not limited to scientific experiments. For two idealized experiments, a distance between these studies quantified by differences in reproducibility rates of the result is a natural measure of how different these two experiments are within the context of replication. Such a distance measure by definition creates a well-defined mapping from experiment space to the space of reproducibility rates, thereby giving a sense of the degree of exactness of replications. When exact replications cannot be performed, the distribution of distance between a study and its non-exact replications provide information about the variability in the replication process. There are a number of difficulties involved in practically defining a distance between studies, however. We describe these in detail after a brief review of the idealized experiment in the next section.

## 3 Idealized experiment and non-exactness of replications

What follows is a modified version of the definition provided in Buzbas et al. (2023, Section 2.1). Assuming some background knowledge *K* on a scientific phenomenon, a scientific theory makes a prediction in principle testable using observables, the data *D*. A scientist formulates a mechanism generating *D* under uncertainty and represents it as a probability model *M* including its assumptions. Given *D*, the scientist is interested in performing inference on some unknown part of *M*. To assess to which extent the desired inference is confirmed by *D*, the scientist uses a fixed and known collection of methods *S* evaluated at *D*. This description captures some key components of studies whose population characteristics can in principle be tested. *S* can be broken down into two components: *S*_*pre*_ and *S*_*post*_. *S*_*pre*_ comprises the set of *scientific* methodological assumptions preceding data collection and procedures implemented to obtain *D. S*_*pre*_ captures the premises underlying the design and execution of an experiment such as experimental paradigms, procedures, instruments, and manipulations. *S*_*post*_ comprises the set of *statistical* methods applied on *D* to obtain the result. *D* can also be broken down into two components: *D*_*v*_ and *D*_*s*_. *D*_*v*_ are data values independently drawn under *M* in each study. *D*_*s*_ is data structure including sample size and sample design. In this paper, we assume that *D*_*v*_ are fixed in each study unless stated otherwise and therefore statements are conditional on a sequence of *D*_*v*_ in these studies. Conceptually, statements unconditional on *D*_*v*_ can be obtained by integrating over appropriate sample spaces, but for this is a tangent to the ideas developed here.

The idealized experiment is the (ordered) tuple: *ξ* := (*K* ≺ *S*_*pre*_ ≺ *M* ≺ *S*_*post*_ ≺ *D*), where ≺ denotes that the component on the left *epistemically precedes* the component on the right^4^. From left to right, the components are mathematically more precisely defined and their properties are better-understood. Each component is conditional on at least on the component immediately to its left and possibly conditional on other components to its left. Serious difficulties arise in quantifying the effect of early components in the tuple, especially *K* and *S*_*pre*_. For example, *K* determines *S*_*pre*_, which may include the types of variables of interest and measurement scales. These in turn restrict the class of reasonable *M* for mechanisms generating the data because only some families of distributions are meaningful under certain types of variables. *M* restricts statistically valid *S*_*post*_ because only some statistical methods are applicable under some classes of models and so on.

We let *r* be a fixed value for result type *R* of interest (which may depend on a user-defined decision criterion). For example, *R* can be the result of an hypothesis test, and *r* might be *choose H*_*o*_. *ξ* induces a definition for the *replication experiment* and the *reproducibility rate* of *ξ* (again, here we provide a modified version of some definitions introduced in Buzbas et al., 2023). An *exact replication experiment ξ*^′^ := (*K*^′^ ≺ *S*_*pre*_ ≺ *M* ≺ *S*_*post*_ ≺ *D*^′^) is equivalent to *ξ* except that it adopts the set

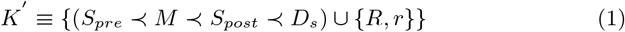

as its background information. This must be so since an epistemic replication assumes knowledge of the model, the methods, and the data structure of the original experiment and aims to replicate them. It does not need to adopt the background knowledge of the original experiment beyond what is needed to repeat it. However, since it is designed with the aim of reproducing an original finding, any replication (exact or not) also needs knowledge of the original result type and the result value to be able to compare its own result to that of the original. It generates random data 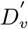 independent of *D*_*v*_. If at least one component differs from *S*_*pre*_, *M, S*_*post*_, *D*_*s*_ or equation 1 does not hold then *ξ*^′^ is a *non-exact replication* of *ξ*. We let *ξ*^(1)^, *ξ*^(2)^, *…, ξ*^(*N*)^, be a sequence of idealized experiments such that all components *S*_*pre*_, *M, S*_*post*_, *D*_*s*_ are identical but the data values are randomly and independently obtained in each experiment. We define the *reproducibility rate* of *r* by

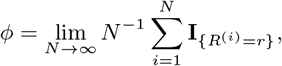

where **I**_{*C*}_ = 1 if *C*, and 0 otherwise. Thus, the reproducibility rate *ϕ* ∈ [0, 1] is a parameter of the sequence of exact replication experiments with respect to *r*. We immediately have 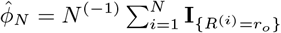 as a natural estimator of *ϕ* since the relative frequency of reproduced results 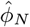 converges to *ϕ* as *N* → ∞ and we are formally comforted to know that lim_*N* →∞_ *P* (*ϕ*_*N*_ = *ϕ*) = 1 by strong law of large numbers. That is, with high probability, the estimated reproducibility rate 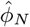 from a sequence of replication experiments gets closer to the true reproducibility rate of the original experiment *ϕ*. Thus, for any given fully specified sequence of exact replications of an experiment up to the unknowns regarding *r*, we have 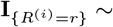 Bernoulli(*ϕ*).

{*R*^(*i*)^ = *r*} is a necessary component to determine whether a specified result is categorically attained. This is straightforward when the statistical inference maps on a categorical decision space. Examples include hypothesis testing and model selection. For other modes of statistical inference, such as point or interval estimation where the natural inference space might be too large to provide practical categories, one needs to be supplied. There have been different approaches to this problem in literature. For example, in Buzbas et al. (2023), we simply assumed that a result is reproduced if interval estimates overlapped in two studies in the context of linear regression. This is unbiased but it decreases precision. This and other methods to measure replication success of a result and their properties need to be carefully evaluated (see Schauer and Hedges, 2021, for some comparative analyses). For point estimation or point prediction, one example is to take the result as reproduced if the point values imply a decision relevant to the scientific inference and not reproduced otherwise (see the example we present in Section 5). Notably, metrics of replication success also impact the reproducibility rate, in addition to experimental components.

Recent replication efforts (e.g., Camerer et al., 2016; Klein et al., 2022, 2018; Open Science Collaboration, 2015) thus far made it clear that assuming that components of studies in the sequence *ξ*^(1)^, *ξ*^(2)^, *…, ξ*^(*N*)^ are identical except the random and independent data is highly optimistic. One reason is practical infeasibility in performing exact replications. A second–and statistically worrying– reason is preference of researchers in choosing different contexts, instruments, procedures, or populations even when it is practically feasible not to do so. Regardless of the reason, *ξ*^(1)^, *ξ*^(2)^, *…, ξ*^(*N*)^ is realized as a sequence of non-exact replications where the definition of an exact replication is no more satisfied^5^. Dispersion of reproducibility rates from non-exact replication studies induces a distribution of reproducibility rates on [0, 1]. Backed up by a properly defined distance, distributions of reproducibility rates can tell us much about differences between studies within fields and between fields. In the ideal case of sampling a sequence of exact replication studies where there is a single true reproducibility rate for a result, this distribution is degenerate. In a sequence of non-exact replication studies, the variability in the distribution of reproducibility rates sampled speaks to the ability of a field to conduct replication experiments. In scientific fields where strict study controls are feasible, we expect the variance of the distribution of *ϕ* to be low, unimodally concentrated around *ϕ*_*o*_ of *ξ*_*o*_. At this end of the spectrum, experimental particle physics comes to mind. Conversely, if strict study controls are not feasible in a field, we expect the variance to be large. At this opposite end of the spectrum, social psychology comes to mind. It might seem that the largest variance occurs when each study in a sequence of studies are different from each other in a way to create a uniformity of reproducibility rates on [0, 1]. This is not the case. To get a feel for this, take Beta(*α, β*) distribution as a reasonable model for *ϕ* ∈ [0, 1]. The worst case of inability in replicating a given study with respect to variance of *ϕ* is realized when *ϕ* ∼ Beta(*α* = *c, β* = *c*) and *c* → 0; that is, when studies with reproducibility rates close to 0 and 1 are conducted at equal frequency. Intuitively, in this situation we are not only unable to repeat a given study faithfully, but we also conduct studies that admit the most-discrepant reproducibility rate with respect to the original study while trying to do so.

## 4 Replication distance

We require that discrepancy between two experiments, captured by a distance, is meaningful in each component *K, S*_*pre*_, *M, S*_*post*_, *D*. Therefore, the distance between two experiments (ideally) is a five-dimensional object. On several occasions where we have introduced this idea, scientists have asked whether the measure can be reduced to a single summary number. Reducing a multidimensional distance to one dimension using a norm such as Euclidean is straightforward algebraically, but the resulting distance will not be interpretable because we know little about the spaces on which these components live. For example, study A can be *d* units apart from both study B and study C, whereas their distance from A has qualitatively different meanings if one differs from A in *S*_*pre*_ and the other in *M*. Thus, our answer is currently no to a unidimensional measure.

To obtain a sensible distance between two experiments that can reasonably be conceived as replications of each other given a result of interest, we impose the following axiom:

At least one component of the same type is *permutable* between two experiments. By permutable, we mean that swapping any component of *ξ*^*o*^ with the corresponding component of *ξ*^′^ is meaningful with respect to the result of interest with well-defined sampling distributions.

Given a fixed component, by the axiom the minimum number of valid permutations is 2. The maximum number of permutations is 2^4^ (if all components result in valid swaps). Conditional on *ξ*_*o*_, *ξ*^′^ we define the reproducibility rate with respect to a fixed component as the mean of reproducibility rates over all permutations. Distance between *ξ*^*o*^ and *ξ*^′^ with respect to the fixed component then is the difference between these averaged reproducibility rates.

More specifically, we let 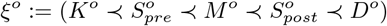 be an original experiment and 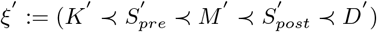 be a replication experiment. Further, we denote the number of permutations for *K, S*_*pre*_, *M, S*_*post*_, *D* by *n*(*K*), *n*(*S*_*pre*_), *n*(*M*), *n*(*S*_*post*_), *n*(*D*), respectively.

We define the distance between *ξ*^*o*^ and *ξ*^′^ with respect to *K* by

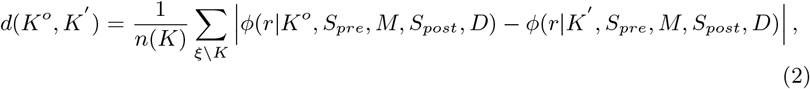

where the sum denotes the permutations over non-fixed components *S*_*pre*_, *M, S*_*post*_, *D*, that is the set *ξ \ K. D* refers to the data structure and not to data values realized in an actual experiment (recall that *D*_*v*_ are fixed for each experiment). The number of relevant permutations scales *d*(*K*^*o*^, *K*^′^) to be in [0, 1], since *ϕ* ∈ [0, 1].

In a fashion similar to equation 2, other distances between *ξ*^*o*^ and *ξ*^′^ with respect *S*_*pre*_, *M, S*_*post*_, *D*, are defined by

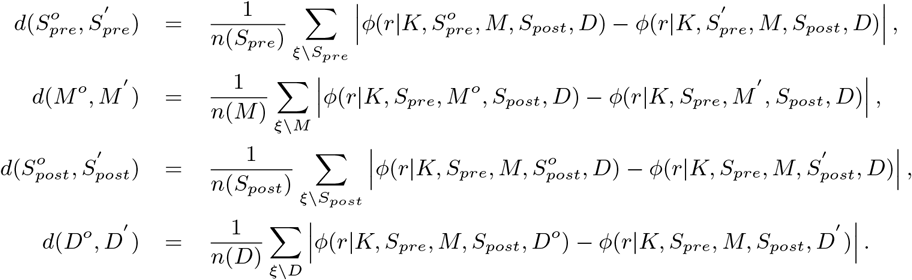

Non-valid permutations require a careful treatment. Here, we assign them reproducibility rate zero by convention by the following reasoning. A non-valid permutation implies that its experiment is not performable and there seems to be (at least) two reasonable ways to handle this case. One way is to assume that the experiment returns Not-A-Number for reproducibility rate and exclude all non-valid permutations from all calculations. However, this reduces the number of cases over which the mean is taken to calculate distances, inflates the reproducibility rate for one component, and consequently decreases the distance unduly. To prevent this effect, by convention we set the reproducibility rate of a non-valid experiment to be zero. The only case that this may cause a problem is when *all* permutations are non-valid, resulting in distance to be zero (by 0-0) and this case is excluded by the axiom which assumes that at least the original two experiments are valid. This formulation yields *n*(*K*) = *n*(*S*_*pre*_) = *n*(*M*) = *n*(*S*_*post*_) = *n*(*D*) permutations. However, to acknowledge the possibility of more advanced metrics that might assign different status to non-valid permutations in order to improve the interpretability of the calculated distances, we keep the notations for the number of permutations in the distance formulas distinct from each other.

We note that using the reproducibility rate, and not the experiments themselves, as a currency makes these distances metric. For example, a distance metric needs to satisfy the condition that the distance between distinct objects is positive. By expression 1, an original experiment and its exact replication are distinct objects. However, for a meaningful replication distance measure, intuitively we desire that the distance between an original experiment and its exact replication is zero so that any positive distance would imply a non-exact replication. While we cannot satisfy this requirement using experiment itself as a currency, we can satisfy it by using reproducibility rate of a result instead, as reproducibility rates from an original experiment and its exact replications are equal.

There are advantages to defining a multi-dimensional distance metric: Since it returns a vector that represents distances across the components of an experiment, two distinct replications that vary in different components would not produce equivalent distances with respect to an original study even if they affect the result (and its reproducibility) in the same way. Of course, it is also possible that, say, the replications differ on two different aspects of the same experimental component (e.g., treatment conditions and dependent measure, both of which belong to *S*_*pre*_) in a way to return the same reproducibility rate. In that case, the distances between each replication and the original would be equivalent. This is not a counter-argument against the internal consistency of our distance measure. It means, given a specific result from an original study and the particular combination of experimental components, these two replications are practically no different from each other and the pre-data methods distance between them is zero with regard to the original. When interpreting these results, it is important to remember that reproducibility rate does not and cannot track an underlying truth independent of the experiment, but rather is prone to reflecting the features of an experiment and what it is capable of capturing. Therefore, two different but compatible replication designs returning a zero (or sufficiently small) distance value with respect to an original experiment is theoretically consistent with the purpose of the metric. It reflects a core assumption that each component of the idealized experiment defines a meaningful categorization and means different replication studies can be effectively conceived that would generate equivalent evidence to test the result in question.

Here we also note that for binary results, a direct consequence of the distance definitions is that distances between two experiments conditional on result *r* and its complement *¬r* are equal to each other by *ϕ*(*r*|*ξ*) + *ϕ*(*¬r*|*ξ*) = 1. That is: |*ϕ*(*r*|*ξ*^*o*^) − *ϕ*(*r*|*ξ*^*′*^)| = |(1 − *ϕ*(*¬r*|*ξ*^*o*^)) − (1 − *ϕ*(*¬r*|*ξ*^*′*^))| = |*ϕ*(*¬r*|*ξ*^*′*^) − *ϕ*(*¬r*|*ξ*^*o*^)|.

Assessing the interchangeability of components, and in fact even defining the measurability of earlier components of an experiment are real conceptual challenges. We give some examples. Theory and technology of scientific studies do not have well-developed tools to work with *K*. If we let *P* be the probability mechanism generating the data, the assumption *P* (*D*|*K, S*_*pre*_, *M, S*_*post*_) = *P* (*D*|*S*_*pre*_, *M, S*_*post*_) must be made to move forward with statistical inference. The left hand side of this equation is *what it is* and the right hand side is *what we operationally use* as scientists and statisticians. The equality holds only when all the available background information about the system relevant to the experiment is captured in *S*_*pre*_, *M, S*_*post*_, *D*. Put differently, given *S*_*pre*_, *M, S*_*post*_, *D*, the data values are independent of *K*. This is a big assumption and making it has significant consequences for our understanding of results of an experiment. *K* emphasizes the fact that the components of the experiment are conditional on what we know about the system before thinking about an experiment. Current approaches to formulating a scientific question from a theoretical scientific model to an experimental model, and ultimately to a statistical model do not seem conducive to quantify *K*.

However, a clear benefit of defining replication distance using reproducibility rate is that we do not need to quantify the components of a replication to be able to calculate its distance from an original experiment. We only need to be able to operationalize the components in the replication as different from the original. That is, as long as we can swap components (say, a *K*^′^ that invokes a different set of instruments than *K*^*o*^) we can compute the distance between the two experiments by calculating reproducibility rates of results under each version. In a philosophical as well as practical sense, this property makes our distance measure real `a la Hacking’s experimental (or entity) realism (Hacking, 1981): If we can manipulate it, it exists.

We see building a mathematical theory of *K*, to formally incorporate it into *P*, and thereby into the likelihood function, as a major challenge in understanding replications and reproducibility. Historically, statistics has not picked up this task, possibly due to lack of theoretical tools to deal with this problem. On the other hand, even when the statistical theory is sufficient to assess an issue, it might be overlooked by scientists. For example, it is well-known that a (simple) random sample from a randomly sampled subpopulation from a population is not equivalent to a (simple) random sample from the population. In many systematic replication studies these samples are assumed to be equivalent. This is a result of science’s drive for generalization of effects, one that is not necessarily supported by statistical theory. On account of these theoretical limitations, for the current proposal of distance provided here, we assume a four-dimensional distance metric instead of five and leave permutations of *K* out of consideration.

Statistical models representing data generating mechanisms in contemporary scientific studies often do not admit closed form solutions. If distances unconditional on data values *D*_*v*_ are desired, they have to be estimated by numerical or simulation-based methods. There is no conceptual challenge in this task and the error in estimates are constrained only by computing power. For example, the distances can be estimated by first estimating the reproducibility rate of each fixed experiment for all experiments using large number of randomly generated data sets *D*_*v*_ and by then Monte Carlo integration. To estimate *ϕ*(*r*|*K, S*_*pre*_, *M, S*_*post*_, *D*), we would simply: 1) Simulate a large number of data sets obeying *K, S*_*pre*_, *D* under model *M*, 2) Analyze it with *S*_*post*_, and 3) Calculate the reproducibility rate as the ratio of results *r* obtained. These estimates have well-known Monte Carlo error bounds.

## 5 A toy example illustrating replication distance

A major use of assessing distance between studies is to inform the design of new studies. Here, we describe a toy example which consists of modified versions of two growth models for wild dugongs (*Dugongs dugon*) to show how distances between two studies are calculated and discuss how the results from distance calculations can be used in practice to design replication studies^6^. Dugong growth models have been used as examples in statistical literature to show the performance of statistical methods in Bayesian (Carlin and Gelfand, 1991; Spiegelhalter et al., 1996) and frequentist contexts (Malouche, 2023).

Dugong growth models predict the length of individuals as a function of age, weight, and more recently other variables involved in age determination such as telomere length (Cherdsukjai et al., 2020). Combined with well-established facts about dugong growth, the relationships between these variables help test life-cycle features of dugongs. In our example here, we use relationships between length, age, and weight to determine whether a study predicts a newly adult dugong to be in the class of ‘certain to breed’ individuals. We have chosen this result because it highlights some properties of distance clearly, not to claim any biological relevance. In fact, distance calculations presented here do not use raw data from actual studies but only simulated data.

Age is measured in half-years, denoted by *x*_1_ ∈ [0, 70], weight in kg *×*10^−2^ ∈ [0.2, 9], denoted by *x*_2_, and response length in meters, denoted by *y* ∈ (1, 3). These boundary values are chosen to be compatible with biology of dugongs from the literature. The maximum age at which dugongs are estimated to reach breeding age is 17 years. Individuals shorter than 2.20 m are unlikely to breed, those longer than 2.50 m are likely to breed, while the status of individuals having body length in (2.20 m, 2.50 m) is uncertain. We take the minimum mean weight to breed at age 17 years as 250 kg and use the estimated model from a study to predict the mean length of 17 years old dugong weighing 250 kg. If the predicted mean length is *>* 2.50 m, an individual is classified as *certain to breed* and otherwise *not certain to breed*. For purposes of result reproducibility we take the result = 1 if the status assigned under the true model matches the status predicted from a replication study, and result = 0 otherwise.

### 5.1 Study 1 specifications

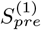 includes predictors log-transformed age and linear weight, and response length. *M* ^(1)^ is a linear regression model following Gauss-Markov assumptions with normally distributed errors, which gives the model

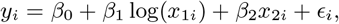

where *ϵ*_*i*_ ∼ *Nor*(0, *tau*), *Cov*(*ϵ*_*i*_, *ϵ*_*j*_) = 0 for all *i≠ j*, and *τ* is the precision parameter. We assume the following sets for true parameter values generating the data: *β*_0_ ∈ (0, 3), *β*_1_, *β*_2_ ∈ (0, 1), and *τ* ∈ [50, 250]. The lower bounds for regression parameters are zero because length is nonnegative and parameters are positive because growth is positive. The upper bound 3 for *β*_0_ is set to cover the maximum length assumption of 3 m and the upperbound 1 for *β*_1_, *β*_2_ are set to simulate data in biologically relevant part of the space. For problems in which no contextual information is available to bound the parameter space, appropriate bounds can be estimated by running a pilot study and regions of parameter space which generate uninteresting data that return only one value (e.g., 0) for the result can be excluded from the analysis.

We set standard deviation of *ϵ* to vary on (0.06, 0.14) based on previous analyses of dugong data (Carlin and Gelfand, 1991). 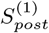 consists of Ordinary Least Squares analysis for the linear regression model (performed by lm function in R (R Core Team, 2013)). Prediction for length is obtained using point estimates of regression parameters with the model evaluated at age 17 years and weight 250 kg. *D*^(1)^ features small low quality samples, with sample size *n* = 20, and bias where age and weight are modified to be 80% and 120% of their actual value respectively (e.g., data processing error).

### 5.2 Study 2 specifications

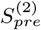 includes predictor age only, and response length. *M* ^(2)^ is a nonlinear regression model with additive errors and is specified as

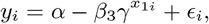

where *ϵ*_*i*_ ∼ *Nor*(0, *τ*), errors are independent of each other and *τ* is the precision parameter. We assume the following sets for true parameter values generating the data: *α* ∈ (1, 3), *β*_3_ ∈ (1, 3), *γ* ∈ (0, 1). The lower bounds for *α* and *β*_3_ are hard bounds implied by the nonlinear growth model form and nonnegative length. The upper bounds *β*_3_ and *α* are set to cover the maximum length assumption of 3 m. Bounds for *γ* are dictated by the desirable property of nonlinear growth model with an asymptote for length at large age. Bounds for *τ* are the same as in study 1. 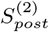 consists of Bayesian analysis with vague priors (nonconjugate): *α* ∼ Nor(0, 10^−6^), *β*_3_ ∼ Nor(0, 10^−6^), *γ* ∼ Unif(0, 1), *τ* ∼ Gamma(10^−3^, 10^−3^). For this nonconjugate model, MCMC (Gibbs Sampler, performed by jags (Plummer et al., 2003)) is used to sample the posterior distribution for each data set generated. For each analysis, an MCMC run consisted of 1 chain, 1000 burn-in steps, and 1000 steps sampled with no thinning. Prediction for length is obtained using the posterior predictive distribution, summing over the posterior samples evaluated at age 17 years and weight 250 kg. *D*^(2)^ features large high quality samples, with sample size *n* = 100 and no errors unaccounted for by the model.

### 5.3. Study specifications in permutations

Sixteen possible study permutations and the reproducibility rates obtained from 10^4^ Monte Carlo simulations for each permutation are given in table 1. Studies 9-12 characterized by 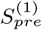, *M* ^(2)^ define non-permutable studies because *M* ^(1)^ requires data on *x*_2_ but 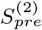 do not provide it. For these permutations, no analysis can be performed and by our adopted convention, the reproducibility rate of the experiment is 0. On the other hand, studies 5-8 characterized by 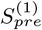, *M* ^(2)^ define permutable studies, the information provided by 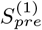 on *x* is simply ignored by *M* ^(2)^.

**Table 1.** All study permutations based on toy example described in section 5. Colored cells show example calculation for model (left), *dist*(*M* ^(1)^, *M* 2), which is the difference between the mean of blue cells and orange cells (right).

| $\xi$ | $S_{pre}$ | $M$ | $S_{post}$ | $D_s$ | $\phi$ |
| --- | --- | --- | --- | --- | --- |
| 1 | 1 | 1 | 1 | 1 | 0.637 |
| 2 | 1 | 1 | 1 | 2 | 0.994 |
| 3 | 1 | 1 | 2 | 1 | 0.637 |
| 4 | 1 | 1 | 2 | 2 | 0.994 |
| 5 | 1 | 2 | 1 | 1 | 0.873 |
| 6 | 1 | 2 | 1 | 2 | 0.944 |
| 7 | 1 | 2 | 2 | 1 | 0.976 |
| 8 | 1 | 2 | 2 | 2 | 0.990 |
| 9 | 2 | 1 | 1 | 1 | 0 |
| 10 | 2 | 1 | 1 | 2 | 0 |
| 11 | 2 | 1 | 2 | 1 | 0 |
| 12 | 2 | 1 | 2 | 2 | 0 |
| 13 | 2 | 2 | 1 | 1 | 0.878 |
| 14 | 2 | 2 | 1 | 2 | 0.950 |
| 15 | 2 | 2 | 2 | 1 | 0.976 |
| 16 | 2 | 2 | 2 | 2 | 0.990 |

| Component 1 | Component 2 | Distance |
| --- | --- | --- |
| $S_{pre}^{(1)}$ | $S_{pre}^{(2)}$ | 0.406 |
| $M_1$ | $M_2$ | 0.539 |
| $S_{post}^{(1)}$ | $S_{post}^{(2)}$ | 0.036 |
| $D^{(1)}$ | $D^{(2)}$ | 0.110 |

Studies 3 and 4, characterized by *M* ^(1)^, 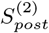 are Bayesian analysis of the linear regression model and they require priors distributions on regression parameters of *M* ^(1)^ to be specified. We assign them vague priors, same priors as in model *M* ^(2)^ for *α, β*_3_: Nor(0, 10^−6^). In this case, vague priors bring very little information into the analysis and they have little effect on the posterior distribution both for *M* ^(1)^ and *M* ^(2)^. However, (*β*_0_, *β*_1_, *β*_2_) are distinct from (*α, β*_3_) as parameters of linear and nonlinear models correspondingly. If informative priors were assigned to (*α, β*_3_) in *M* ^(2)^ of study 2 specification, we would be amiss to assign the same priors to (*β*_0_, *β*_1_, *β*_2_). If this were the case, our first attempt would be to look for transforms of the priors on (*α, β*_3_) to priors on (*β*_0_, *β*_1_, *β*_2_). If no transforms are found, we would label studies 3 and 4 as non-permutable.

The reproducibility rates of studies 5 and 6 are equivalent to studies 13 and 14 respectively in this problem. This is due to the fact that for 5 and 6 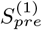 provides *x*_1_, *x*_2_ as predictors but *M* ^(2)^ uses only *x*_1_, resulting in the same study permutations as in 13 and 14 respectively. A consequence is that the net effect of studies 5, 6, 13, and 14 on distance of a varying component is approximately zero for that component. These studies pair model *M* ^(2)^ and 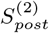, and the estimation problem is a Nonlinear Least Squares problem. Well-established methods to the Nonlinear Least Squares problem are numerical gradient-based methods. A standard is the Levenberg–Marquardt algorithm, which we use through R package minpack.lm. These numerical methods, although state of the art, have computational limitations. For example, for certain parameter values which yield near-singular matrices, methods may not be able to calculate gradients accurately and they return NAs for parameter estimates. In our case, few parameters resulted in this condition due to well-constrained parameter space. We excluded the cases where NAs are returned from the calculations and this choice had little effect on the reproducibility rates. However, it is easy to imagine problems with large parameter spaces where the frequency of NAs may be high enough (implying that the method is not appropriate for the model) and the reproducibility rates are substantially affected by this shortcoming.

### 5.4 Interpretation of distances

Based on the reproducibility rates given in Table 1 (left) for study permutations, the distances between experiments for each component are calculated by averaging over all other components (right). Let us assume that we want to interpret these distances with a goal to inform new replication studies. The idea is to assess the sensitivity of each component with respect to the result. Sensitive components would require more caution and higher fidelity to the original in future replication studies than nonsensitive components.

Pre-data methods are far from each other (0.406) with respect to the given result. This is mainly due to 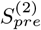, *M*_1_ yielding non-permutable experiments (since lack of weight as a predictor in *S*_*pre*_ cannot accommodate *M*_1_ which uses weight) and consequently returning reproducibility rate zero. The inclusion/exclusion of weight as a predictor also yields a large distance between models (0.539). It might seem that 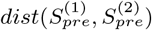 and *dist*(*M* ^(1)^, *M* ^(2)^) would be highly correlated for all experiments, but this is not necessarily the case. The reason why they are correlated for this example is that the predictor (weight) omitted in 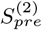 has a nonzero effect on the response (length) and including this predictor in the model results in large reproducibility rate in 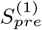, *M* ^(1)^ permutations. Consequently, the gap in reproducibility rates with 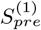, *M* ^(1)^ and zero cases is large, contributing to the distance between models as well. Thus, when designing replication studies, it would be to our advantage to choose an *S*_*pre*_ that includes all predictor variables that are believed to have a large effect in the set of models of interest, or alternatively, only to consider models that can be accommodated by a fixed *S*_*pre*_, so that the cases that return zero reproducibility rate are eliminated.

The distance in post-data methods is negligible (0.036). This is expected. The Least Squares (ordinary and nonlinear) regression based on the frequentist approach, and the posterior predictive based on the Bayesian approach using *vague priors* are well-behaved methods under the models considered here. The error variance in the data generating mechanism is small, models have few parameters, and even the non-linear relationship is simple, making statistical inference straightforward. Thus, in a new replication experiment we can afford to choose our desired method of analysis with the knowledge that it would not have a substantial effect on the results. A word of caution here is that vague priors are an important part of this conclusion. Informative priors may increase 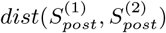 very easily, of course.

The distance between *D*^(1)^ and *D*^(2)^ is very small (0.110). This is unexpected because the data quality in *D*^(1)^ (small sample, bias in predictors) is considerably lower than that in *D*^(2)^ (large sample, unbiased predictors). First, this result informs us that before performing a new experiment, a sensible approach would be to study the effect by designing further simulation studies, for example trying different biases in predictors to assess their effects. Second, if we are ready to accept *dist*(*D*^(1)^, *D*^(2)^) = 0.110 at the face value, the implication is that the effect of the difference in data quality *with respect to the conditioned result* as measured by reproducibility rate is minimal. However, we emphasize that this result does not imply that the effect of data quality will be necessarily similar for other result types.

## 6 Minimum Viable Experiment to replicate and planned modifications

Recently, a large number of systematic replications of various results have been conducted. Unfortunately, most of these purported replication studies differ from the original study in at least one–and often more than one– component *S*_*pre*_, *M, S*_*post*_, *D*. Therefore, they are at best non-exact replications of an original study; worse yet, some don’t even qualify as proper replications due to all components being non-permutable or conditioning on a different result type. We have established that reproducibility rate of results in haphazard non-exact replication studies are highly variable (Buzbas et al., 2023). Since non-exact replications are practically inevitable, we need to find a way to perform them in a controlled and systematic manner and specify their aim. To that end, we have created the concept of *Minimum Viable Experiment* to replicate (MVE, Devezer and Buzbas, 2022).

MVE is an elementary experiment characterized exclusively by a minimum number of indispensable parameters needed to generate the result of interest. Complementing MVE, the sequence of studies performed to identify the MVE aim *not to replicate an original study*, but consist of a sequence of *purposefully planned modification studies* by systematically swapping some of the components *S*_*pre*_, *M, S*_*post*_, *D*, of the previous study in the sequence and aims to iteratively assess the effect of these changes on the result of interest. Modifications that do not yield the desired result are discarded and modifications that yield or improve the desired result are maintained. The ultimate goal is to allow the sequence to evolve in a way to help identify the minimal conditions and variables in a study which make a desired result observable. We say that such a study is replication-ready in the sense that MVE is simple enough to allow for near-exact replications. If the MVE can be identified, the reproducibility rate of its result can be obtained with high precision.

There are some key differences between an exact replication study and a purposefully planned modification study with respect to an original study (see Figure 1 for a comparison). An exact replication receives its background information (*K*^′^, orange text) from all components of the original study because the replication needs to know each experiment to replicate them, as well as the result to check whether the result is reproduced. Then, *K*^′^ feeds all the components of the replication experiment (exact replication, orange arrows). A purposefully planned modification gets the background information from the original experiment (*K*^′^, orange) and combines it with additional background information (*K*_+_, teal) to obtain its background information. *K*^′^ and *K*_+_ then feed all the components of the planned modification experiment together. *K*^+^ informs which modifications are to be made in which components of the planned modification experiment so that their effects on the result of the experiment can be assessed. Thus, *K*^+^ iteratively improves upon *K*^′^.

**Figure 1.**
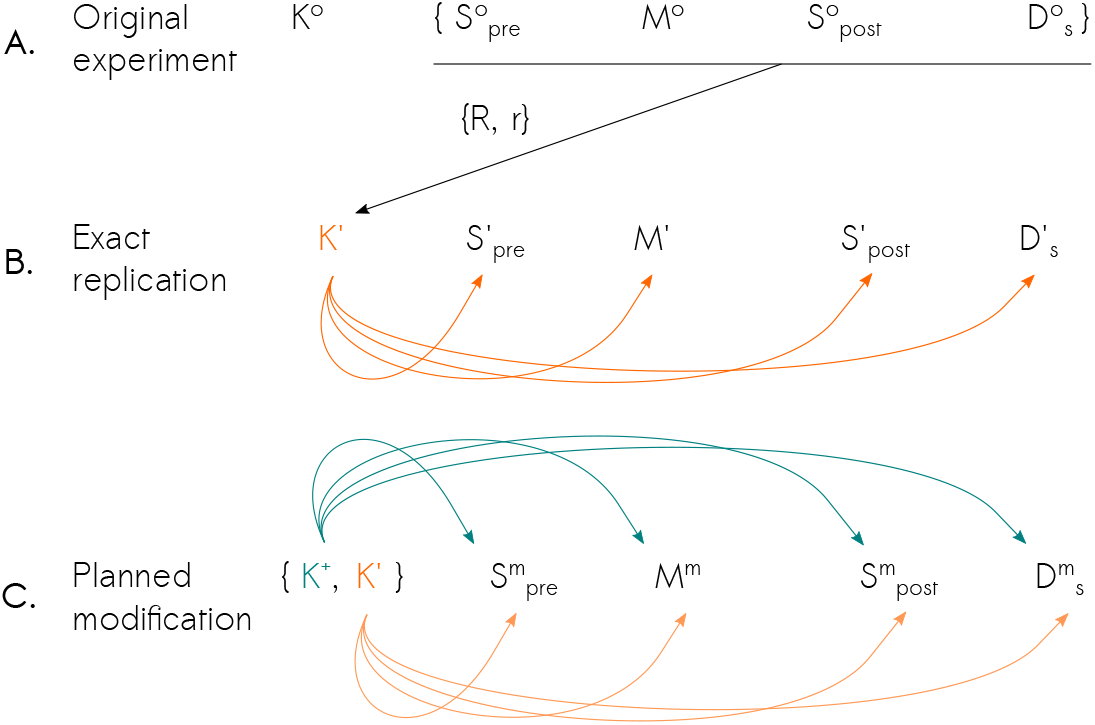
(A) Original experiment, (B) Exact replication’s *K* ^′^ uses all components of the original experiment exactly (except *K*^*o*^), the result type (*R*) and value *r*, (C) Planned modification takes the same information from the original experiment equivalent to *K*^′^ and some external background information *K*^+^, and creates *K*^*m*^ to define its own components.

One interesting way to think about planned modifications is as experimental designs set up over a sequence of idealized experiments. One of the most fundamental properties of scientific experiments guided by statistical experimental design is that the only difference between control and treatment groups is the controlled manipulation of the predictor variables to assess its effect on the response variable of interest. In planned modification studies, the (predictor) variables are akin to components of the idealized experiment, and the (response) variable is akin to the result *r* of interest. Thus each step in the sequence of planned modifications is as if we perform a new treatment based on the (control) experiment given by the previous experiment in the sequence. This systematic approach is also in stark contrast with non-exact replications where modifications are haphazard and often simultaneous.

Figure 2 contrasts the strategies adopted by replication studies, their realizations, and strategies adopted by the MVE schematically. The geometric shape denotes the model’s size (higher order polygons denote larger models), the color (orange/blue) denotes one of two statistical methods considered for analysis, and the shade denotes sample size (darker shades denote larger sample sizes). An example of the two most distant cases is given to the right of the figure: The triangle represents the simplest model considered, using one type of analysis method, and having the smallest sample size (lightest shade). The dodecagon represents the most complex model considered, using the other type of method to analyze the data, and having the largest sample size (darkest shade). Based on an original study *ξ*^(*o*)^, a sequence of replications *ξ*^(1)^, *…, ξ*^(*t*)^ aim to replicate *ξ*^(*o*)^ exactly (A), but instead typically achieve a sequence of uncontrolled non-exact studies (B) jeopardizing the assumptions of replications. The sequence of studies targeting MVE modify the study components in a controlled manner, one component at each step, and test the relevance of modification with the goal of identifying the MVE (C).

**Figure 2.**
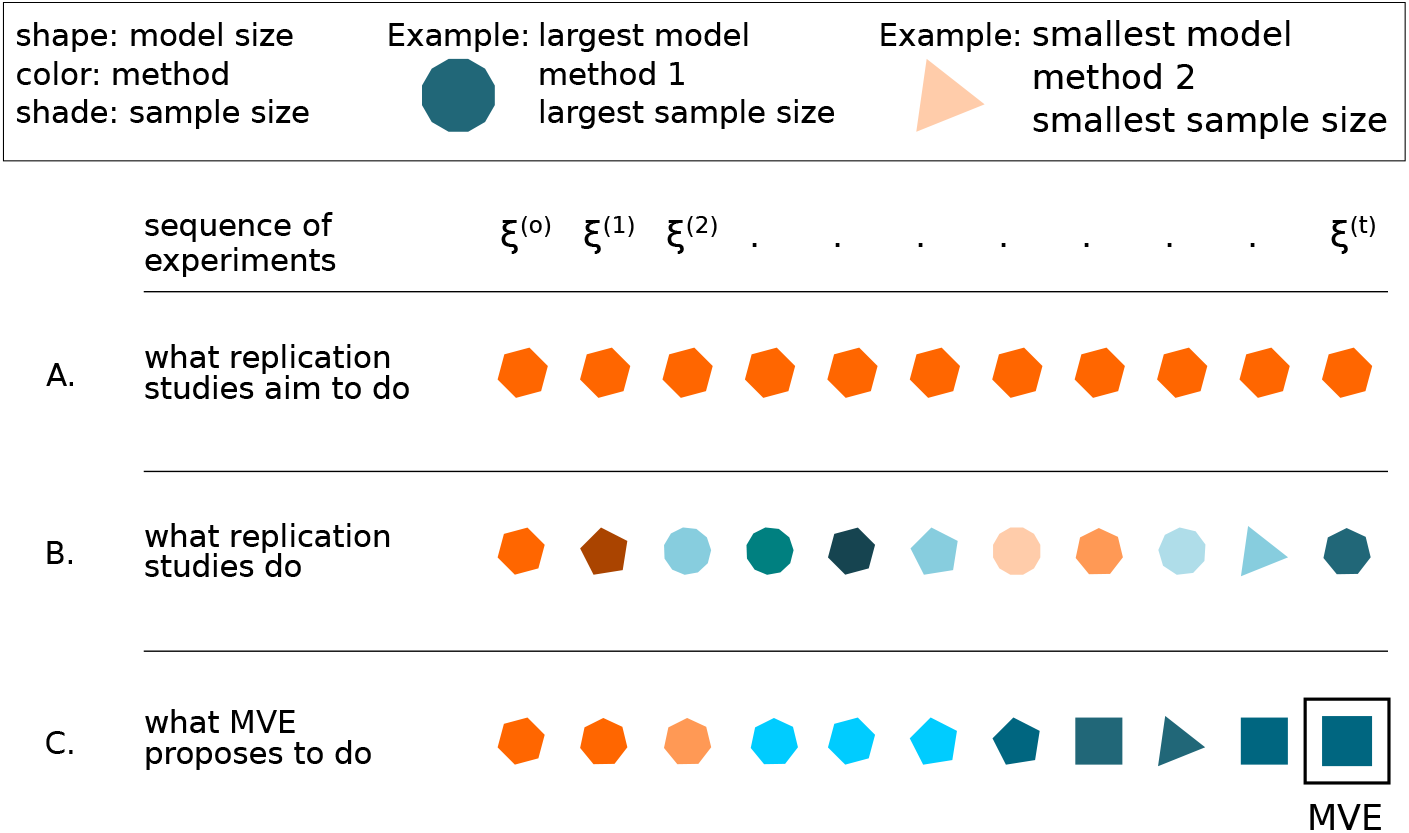
Contrast between the goal of replication studies (A), their currently conducted versions (B), and the evolution to a Minimum Viable Experiment to replicate (C) in a sequence of studies.

The distance described in Section 4 is useful in assessing the differences between planned modifications through a sequence of studies. The application of distance equation 2 (and four relevant equations after) at a given step *t* in the sequence gives 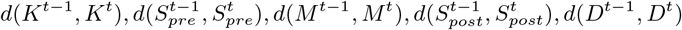.In a sequence of *m* purposefully modified studies, we get *m* − 1 5-dimensional distances, showing the modifications made in each component of an experiment at every step of the sequence and its effects on reproducibility rate of the result of interest. We expect to start with large fluctuations in distances in all dimensions at early steps of the sequence because not much is known about the MVE at exploration stages of a phenomenon. Planned modifications work as planned if they eliminate unnecessary variables and model components (and include the necessary ones), and improve methods to obtain the specific result. Thus we expect that if planned modifications work as intended, fluctuations in distances should decrease to constants in all dimensions as we approach to MVE. The rate of approach to these constants will depend on how sensitive the result is to changes in components, as well as how often the planned modification is applied for an component in the experiment. At an acceptable MVE, that is when a study is replication-ready, all distances will be zero (provided we know the result of the original experiment). There is a lot of room for strengthening the theoretical connections between distance, planned modification, and replication-readiness to guide study choice and design in scientific practice.

## 7 It is about science!

The ideas presented in this paper are about formal ways of incorporating (generally) meta-notions into statistical theory with a specific focus on replicability and reproducibility. We believe that early components of a scientific study (e.g., *K, S*_*pre*_) represent more serious challenges than the later components (e.g., *M, S*_*post*_, *D*). The reason is that early components have not been studied sufficiently whereas statisticians have worked on the latter components extensively.

How does this perspective, complementary to Taper et al. (2022) connect to philosophical approaches to statistics including evidential statistics?

Understanding the nature of evidence when moving from subject-matter-specific scientific models into–largely constrained–experimental models, and then finally to statistical models ready for inference seems to be the biggest challenge for accumulating scientific knowledge. This passage among conceptual models has not been worked out satisfactorily (although see a conceptual attempt we offer in Devezer and Buzbas, 2023). In the context, we believe that the natural approach for building new theoretical ground for connections between scientific inference via statistical inference is to start with a *pure likelihood-based* approach at the early stages of development and after some basics are established extend the framework adhering to *evidential approach*. Our justifications for this preference are as follows.

- Likelihood functions are excellent tools in statistical inference because they solely rely on mechanism generating the data, broadly construed. They are also sub-class of evidence functions and compatible with evidential theory.
- Likelihood functions circumvent limitations and inconsistencies of frequentist statistics.
- Even if one wants to take a Bayesian approach –not necessarily philosophically, but purely due to computational advantages that it provides in simulation-based methods such as Monte Carlo– one has to first work with valid likelihood functions. In addition, likelihood functions would avoid complications due to working with priors in early theory-building for meta-notions.
- The scope of problems imposed by meta-notions are conceptually broader than problems that philosophy of statistics often tackles, which almost always assumes that models hold and then looks forward to develop and evaluate statistical methods^7^. Incorporating meta-notions such as replicability and reproducibility into a philosophy of statistics is rather interested in representing these notions in the models appropriately in the first place. In this sense, evidential theory currently seems the only approach that can join the two functions seamlessly. That is, evidential theory is certainly comfortable to work with *M, S*_*post*_, *D* and promises to be flexible enough to work with objects such as *K* and *S*_*pre*_.

There is no question, however, that trying to incorporate the real life imperfections of doing science into a formal framework is a daunting task. For connecting models and comparing experiments, a valid distance that describes the relationship between *scientific* studies (with all their components *K, S*_*pre*_, *M, S*_*post*_, *D*) seems a necessary first quantity ^8^. We have described an example of such a distance in this paper based on reproducibility rates as a currency (which are based on likelihoods). The progress to MVE, which emphasizes the evolution of evidence as we modify studies also needs this distance as a measure. A evidential approach with its clear focus on evidence functions, especially the class of likelihood functions is naturally suited to clarify this problem.

This preference does not mean that we have no difficulties in representing meta-notions under the evidential theory, however. Using replications and reproducibility as a running example, it is currently not clear whether some of the desiderata in Taper et al. (2022) can be satisfied for some components of an idealized experiment. Specifically:

- It seems challenging to restrict *K* and *S*_*pre*_ so that the induced *M* creates evidence that is continuous function of data (desiderata, D2).
- By its very nature *K* is likely to clash with desiderata D7. (It is also interesting to note that even in well-known cases where statistical theory exists for invariant estimators, misapplications are common).
- Open-world/closed-world problem about the set of *M* has no current solution. If one is invented, it might be incompatible with desiderata D8.

To round up our perspective, we offer a slightly revised version of the question raised by Royall (2004) and reiterated in the call for this special issue by Taper et al. (2022). We believe that the crux of problem in philosophical and theoretical approaches to statistics is the failure in addressing: “When does a given set of observations support one *scientific* hypothesis over another?”

## Author contributions

Conceptualization, E.B. and B.D.; methodology, E.B. and B.D.; writing—original draft preparation, E.B. and B.D. All authors have read and agreed to the published version of the manuscript.

## Footnotes

1 Many data-specific challenges have been tackled in application areas of biometrics, technometrics, and psychometrics. For example, specialized methods for spatioemporal data, survey data, genetic data, and directional data have been invented. However, most of these methodological solutions are not necessarily related to the meta-notions that we discuss.

2 These terms have been defined in conflicting ways in the literature. For internal consistency within our research program, we abide by the definitions provided in Baumgaertner et al. (2019) and Buzbas et al. (2023).

3 Here we rely on the theoretical framework we have been building over the last few years in Baumgaertner et al. (2019); Buzbas et al. (2023); Devezer et al. (2019, 2021).

4 In Buzbas et al. (2023), we have used the notation (*K, S*_*pre*_, *M, S*_*post*_, *D*), and the order is implied by definition of “tuple”. However, it has largely been overlooked in metascience literature and is essential for valid inference.

5 Strong Law of Large Numbers show that we could sensibly talk about a *mean reproducibility rate* 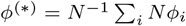 with the corresponding estimate 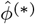 still maintaining the desirable convergence properties, but *ϕ*^(*)^ is challenging to interpret.

6 Computer code used to generate results in this section is available at https://github.com/Devezer-Buzbas/Distance_Between_Experiments.

7 This is not to say that the idea of measuring and comparing experiments has been completely neglected by theoretical statisticians (see for example, Blackwell, 1951, 1953; Lindley, 1956). However, in this line of literature an experiment is viewed as a sampling procedure; that is, a well-defined, strictly statistical entity as opposed to a blurry, imperfect scientific one.

8 The distance measure advanced here is not (yet) a fully developed tool ready for practitioner use. We present it in its early phases of theoretical development to illustrate the fruitfulness of such a theoretical approach. Several versions of the same metric can be advanced based on the information provided here. Their properties need to be well understood before the best candidate can be offered for wide-scale implementation as a ready-to-use tool.

